# Using the chicken embryo as an *in vivo* model to revive viable but non-culturable (VBNC) pathogens

**DOI:** 10.1101/2025.03.21.644564

**Authors:** Filipe Carvalho, Iana Hemery, Catherine Schouler, Alessandro Pagliuso

## Abstract

The chicken embryo has emerged as a popular *in vivo* model with increasing application in biomedical research, due to its simplicity, affordability, and adaptability in the study of various biological phenomena. This model has been used to investigate microbial pathogenicity, and is becoming a useful tool to study bacterial dormancy. The viable but non-culturable (VBNC) state is a dormant state in which bacteria become metabolically quiescent and resistant to cultivation to preserve their viability in harsh environments. Under favorable conditions, VBNC bacteria can “wake up” back into a metabolically active and culturable state. Bacterial pathogens that switch to a VBNC state, such as the foodborne listeriosis-causing *Listeria monocytogenes*, are a public health concern, as they elude detection by conventional growth-dependent methods and can recover their virulence upon revival. This urges a better understanding of the conditions and mechanisms driving the revival of VBNC pathogens. The method presented here showcases the chicken embryo as an efficient *in vivo* model to revive VBNC *L. monocytogenes* back into a culturable status. Where *in vitro* revival attempts, largely based on nutritional replenishing, were unproductive, this protocol succeeds in promoting the reactivation of cell wall-deficient VBNC forms of *L. monocytogenes* generated by starvation in mineral water. Importantly, our results underline the requirement of the embryo for the revival of VBNC *L. monocytogenes*, indicating an important role of embryo-associated factors in this process. Other potential uses for this method include the screening and identification of bacterial factors implicated in the mechanisms of VBNC state revival. This model can thus provide insight into the molecular workings of bacterial dormancy, whose knowledge is critical to reduce the public health risks entailed by undetectable pathogens.

**SUMMARY:** This method showcases the chicken embryo as a simple and cost-effective *in vivo* model to revive the bacterial pathogen *L. monocytogenes* from a viable but non-culturable (VBNC) state, and with potential further uses in the understanding of bacterial dormancy mechanisms.

## INTRODUCTION

In the search for alternative *in vivo* research models, with reduced associated costs, logistical and ethical considerations, the chicken embryo emerged and quickly became one of the most applicable, manageable and reproducible *in vivo* vertebrate model systems. Compared to other animal models, such as rodents and rabbits, fertilized chicken eggs are inexpensive to obtain and require no complex housing logistics for their development. Moreover, the chicken egg size enables the handling of numerous embryos in parallel, supporting a robust number of experimental test conditions/groups and replicates. The short embryogenesis duration (21 days) and the simplicity to access and observe the embryo and associated structures at any point of the developmental period make this a useful model in a variety of fields, such as developmental biology (e.g. heart and brain formation)^1, 2^ and pharmacology (testing drug activity, delivery and toxicity)^3–5^. In addition, the immature immune system of the chicken embryo makes it a suitable system for immune-based studies and cancer research-related approaches^6^. Importantly, because of the embryonic nature of this model, which only acquires a mature nociceptive system by developmental day 13–14^7^, research applications conducted within this timeframe are not constrained by legal and ethical concerns^6^.

The chicken embryo model has also been widely used to study the pathogenicity of microbes causing disease in humans and other mammals. Indeed, numerous studies have explored and validated this model to investigate the virulence of protozoan (e.g. *Neospora caninum, Eimeria tenella, Cryptosporidium* spp.)^8–10^, fungal (e.g. Candida albicans, Aspergillus fumigatus)^11, 12^, and bacterial species (e.g. *Enterococcus spp*., *Salmonella enterica, Francisella spp*., *Campylobacter jejuni, Clostridium perfringens, Listeria monocytogenes, Neisseria gonorrhoeae, Staphylococcus aureus*)^9, 13–21^, as well as to test the therapeutic effect of antimicrobial compounds^16, 22^.

Microorganisms, like the ones mentioned above, are often exposed to stressful stimuli in their environment(s), and have therefore evolved stress-coping strategies to endure through potentially harmful/lethal situations. Some bacterial species can produce highly resilient and metabolically dormant structures called endospores, which preserve the cellular and genetic integrity under environmental constraints. If favorable environmental conditions are gathered, endospores can regenerate into viable active cells via germination^23^. Non-sporulating bacteria, however, may enter into an alternative metabolically dormant state, called “viable but non-culturable” (VBNC), whose main phenotypic trait is the loss of culturability in routine growth media^24^. Given that a large part of the >100 bacterial species reported to enter a VBNC state is pathogenic to humans and other animals^25^, the failure of conventional growth-based methods to detect VBNC pathogens is a concerning public health issue. In addition, VBNC pathogens may “revive” back into a metabolically active state and regain their virulence^24, 25^. The environmental cues and the molecular and physiological mechanisms driving this revival process are not yet well understood, and may vary with the microbial species and the VBNC state-inducing stress(es).

Researchers have taken advantage of the particularities of the chicken embryo model to investigate the *in vivo* revival capacity of bacterial pathogens in a VBNC state. Human-derived *C. jejuni* isolates, driven into a VBNC state by nutritional deprivation in water, recovered their

culturability and virulence in human cells after passage in embryonated chicken eggs^26^. Similarly successful attempts to revert the VBNC state were also reported for other pathogens, such as *Edwardsiella tarda*^27^, *Legionella pneumophila*^28^, and *L. monocytogenes*^29^.

We have recently reported that when *L. monocytogenes* is driven into a VBNC state by starvation in mineral water, it shifts from a rod-shaped to a coccoid cell. We revealed that this morphological transformation is caused by cumulative damages to the cell wall leading into its complete shedding by the bacterium, which then becomes a cell wall-deficient spherical cell form^30^. Our unsuccessful attempts to revive these wall-less VBNC *L. monocytogenes* forms *in vitro*, using nutrient replenishing approaches, led us to investigate their potential rescue *in vivo*. Given its promising use with *L. monocytogenes*^29^, we selected the chicken embryo model for this task. Our results confirmed the capacity of the embryonated chicken egg system to promote the restoration of cell wall-deficient VBNC *L. monocytogenes* back to an active culturable state^30^.

Here, we provide a detailed protocol describing the preparation and monitorization of chicken eggs and VBNC bacteria, the inoculation of eggs, the processing of the embryonated and non-embryonated eggs, and the scoring of the culturable bacterial burden to assess VBNC cell revival efficiency. Our results reaffirm the chicken embryo as a simple, cost-effective and suitable model to understand the mechanisms governing different aspects of microbial life, such as bacterial dormancy. This *in vivo* system can be further explored to investigate the contribution of individual bacterial genes in the resurrection process.

## PROTOCOL

This protocol follows the applicable institutional, French (Decree no. 2013-118) and European (Directive 2010/63/EU) guidelines for animal use in research. Furthermore, under French law (Decree no. 2020-274), this protocol is not concerned by ethical restrictions because all experimentation with chicken embryos is performed and completed before the last third period of embryogenesis (i.e. before day 14).

### 1. Preparation of VBNC bacteria

1.1. Streak bacterial strains from glycerol stocks (stored at −80 °C) onto brain heart infusion (BHI) agar media, using inoculation loops. Incubate overnight at 37 °C to obtain isolated colonies.

NOTE: If required, supplement the agar media with antibiotic(s) for which the strain(s) possess(es) resistance (either naturally or via genetic modification).

1.2. For each strain, prepare 2–3 tubes with 5 mL of BHI broth and use an inoculation loop to inoculate with 2–3 colonies. Incubate the cultures overnight at 37 °C with agitation (200 rpm) to grow bacteria until the stationary phase.

NOTE: Biological replicates of VBNC bacterial suspensions are prepared from independently grown bacterial cultures.

1.3. Dilute each culture 1:10 in BHI broth and measure the optical density in a spectrophotometer at a wavelength of 600 nm (OD_600_).

NOTE: The OD_600_ value of optimally grown stationary phase cultures of *Listeria monocytogenes* ranges between 2 and 4. This equates to a bacterial concentration of 2–4 × 10^9^ colony forming units (CFU)/mL.

1.4. Pellet 1 mL of bacterial culture in a 1.5-mL microtube by centrifugation at 6000 × *g* for 2 min. Aspirate the supernatant using a vacuum pump. Resuspend the cell pellet in 1 mL of autoclaved and filtered mineral water.

1.5. Wash the bacteria thoroughly by repeating step 1.4 three times.

NOTE: These washes are important to completely remove any trace of culture medium nutrients that would, otherwise, delay the formation of VBNC bacteria.

1.6. Prepare bacterial suspensions at a starting concentration of 10^6^ CFU/mL by adding 30 µL of washed bacteria to T25 cell culture flasks containing 30 mL of mineral water.

1.7. Mix the suspensions using a serological pipette. Store them at room temperature, under static (flask upright) and dim lighting conditions.

### 2. Monitoring the formation of VBNC bacteria

2.1. Determine the culturable population in the bacterial suspensions on the first day and then on a weekly basis, as follows:

2.1.1. Prepare 10-fold serial dilutions of the VBNC bacterial suspension in mineral water (from 10^−1^ to 10^−3^ dilutions). Plate 100 µL of undiluted suspension and of each dilution, in duplicate, on BHI agar and incubate overnight at 37 °C.

NOTE: When the culturability value approaches <1 CFU/mL, a larger volume (0.5–2 mL) of the undiluted suspension should be plated to confirm the presence/absence of culturable bacteria.

2.1.2. Calculate the concentration of culturable bacteria (CFU/mL) as follows: (average number of colonies × sample dilution factor) / plated sample volume (in mL).

2.2. Determine the viable population in the bacterial suspensions on the first day and then on a weekly basis, as follows:

2.2.1. Incubate a sample of the bacterial suspension with the viability dye 5(6)-carboxyfluorescein diacetate (CFDA) at a final concentration of 30 µM, for 30 min in the dark. Prepare also a sample without CFDA (unstained control), and a sample treated at 95 °C for 30 min before incubation with CFDA (non-viable/dead cell control).

2.2.2. Analyze the samples in a flow cytometer equipped with a 488 nm excitation laser and a 520 nm emission detector for CFDA fluorescence. Use the unstained and non-viable/dead cell control samples to distinguish the viable (CFDA-positive) from the non-viable (CFDA-negative) populations.

2.2.3. Calculate the concentration of viable bacteria (cells/mL) as follows: average number of CFDA-positive events in acquired sample / acquired sample volume (in mL).

NOTE: Depending on the cytometer model, cell quantification may require the use of flow cytometry counting beads.

## 3. Incubation of eggs

3.1. Upon reception from the supplier, allow the fertilized eggs to accommodate at room temperature until the next day.

3.2. In the meantime, fill the water reservoir of the egg incubator with deionized water and turn the power on. Set the temperature to 37.7 °C and the maximum relative humidity to 47%.

3.3. Transfer the eggs into the incubator to initiate embryogenesis. Position them on the incubator tray(s) with the air pocket facing up (i.e. pointy end facing down).

NOTE: In our hands, eggs are incubated at a relative humidity range of 38–42%, with no visible impact in the expected embryogenesis rate and success.

3.4. Incubate the eggs for 6 days. After 4 days, live embryonated eggs can already be distinguished from dead or non-embryonated eggs by candling (Figure 1).

**Figure 1:**
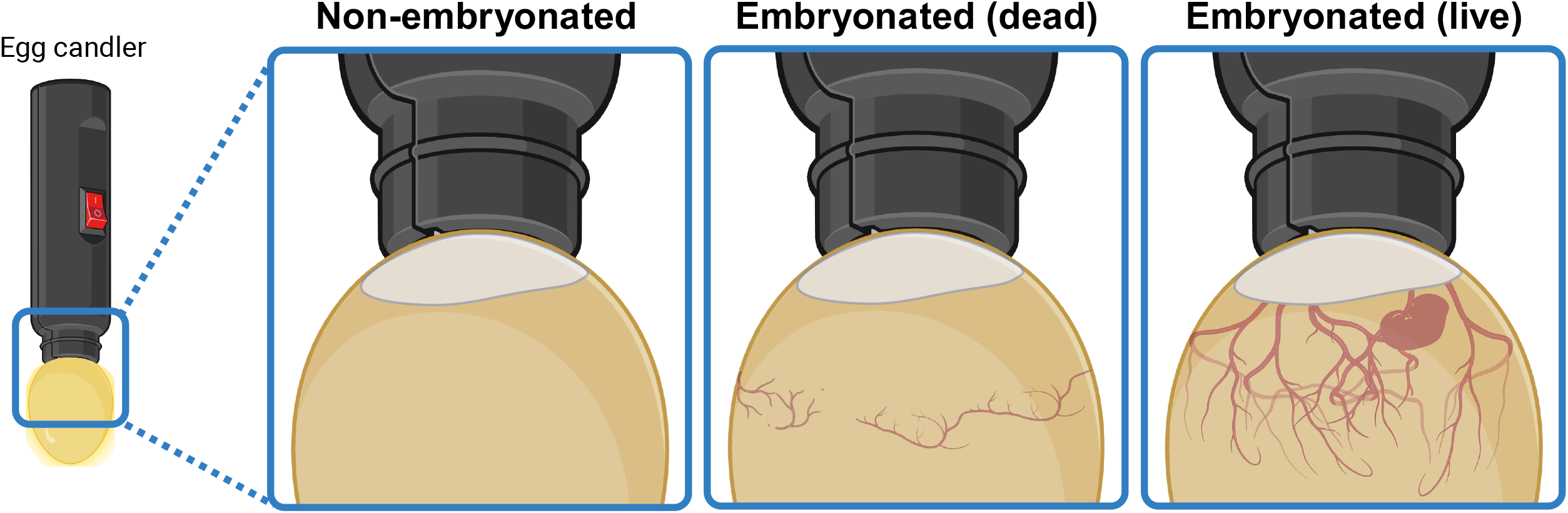
Distinction of non-embryonated, dead embryonated, and live embryonated chicken eggs by candling. Transillumination of chicken eggs through the air pocket end allows to visually determine the embryogenesis status. After 3 or 4 days of incubation, live embryonated eggs display a network of blood vessels expanding downwards from the air pocket. This network appears collapsed and disorganized in dead embryonated eggs, and is completely absent in eggs that failed to start embryogenesis. If visible, a live embryo can display twitching movements. Created in BioRender.com.

### 4. Preparation of eggs for inoculation

4.1. On the day of inoculation, candle the eggs to determine the available number of viable embryonated and non-embryonated eggs. Identify the non-embryonated eggs (about 10–20% of total number) and discard any eggs with compromised embryogenesis.

4.2. Mark the injection spot on the egg shell at 2–5 mm above the air pocket border (Figure 2A). Disinfect this area by rubbing with a tissue paper soaked with 70% (v/v) ethanol.

**Figure 2:**
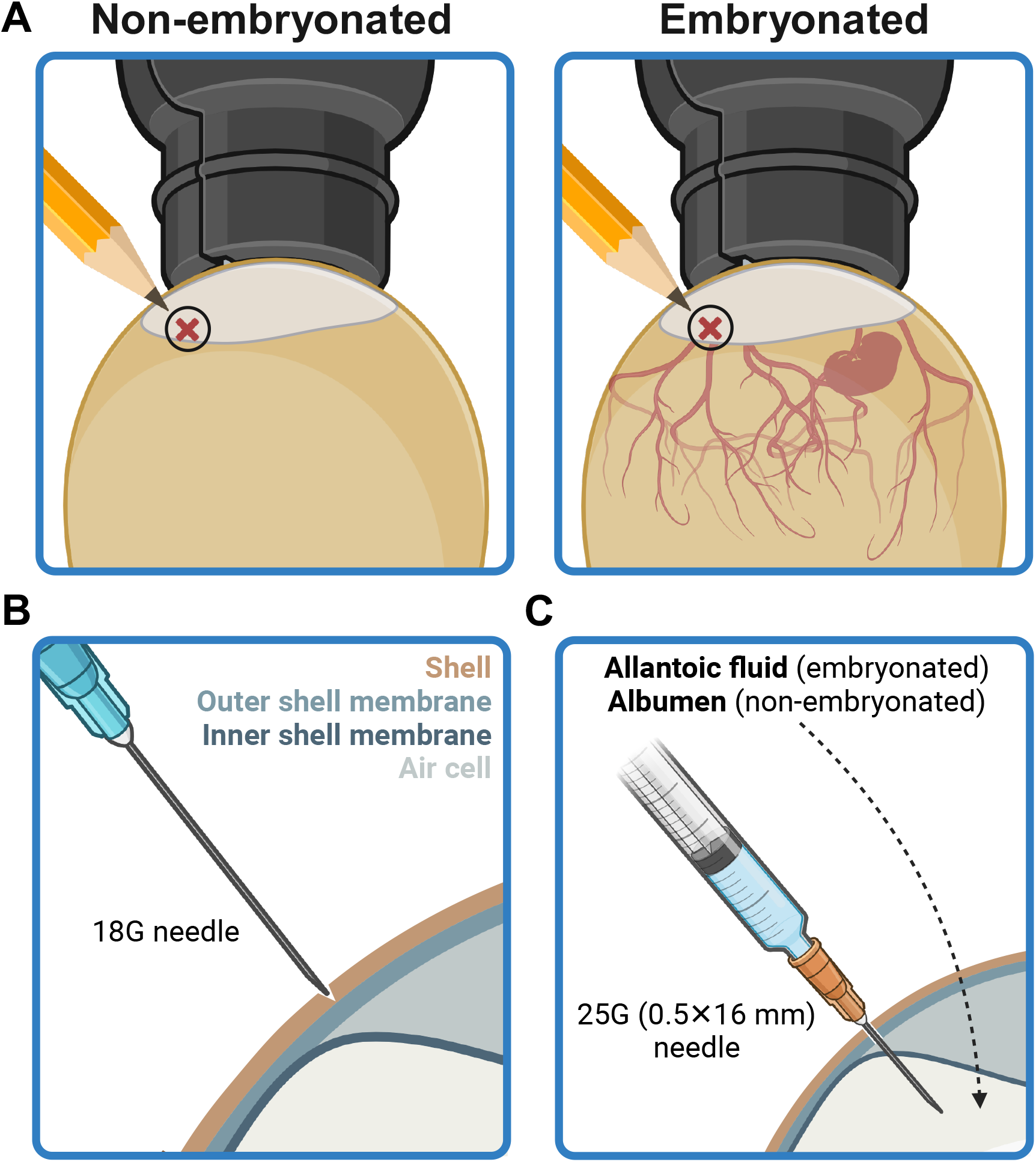
Preparation of the egg for inoculation. **(A)** The injection site is marked at a position 2–5 mm above the air pocket border, as seen by candling. **(B)** The shell at the injection site is puncture dented with the help of the tip of an 18G needle, without piercing the outer shell membrane. **(C)** The inoculum is delivered through a 25G (0.5×16 mm) needle into the allantoic fluid, in embryonated eggs, or the albumen, in non-embryonated eggs. Created in BioRender.com.

4.3. Using an 18G needle, carefully make a puncture dent on the shell at the injection spot, without piercing through the shell membrane (Figure 2B). Disinfect the injection spot with a tissue paper soaked with 70% (v/v) ethanol.

### 5. Inoculation of eggs

5.1. Plug a 25G (0.5×16 mm) needle to a 1-mL syringe and fill it with VBNC bacterial suspension. If necessary, remove air bubbles stuck to the inner wall of the syringe by flicking it with a finger. NOTE: Optionally, inoculation with mineral water alone or a suspension of *L. monocytogenes* (5×10^3^ CFU/mL) grown overnight in BHI broth can be used as control conditions for sterility and growth derived from culturable bacteria, respectively.

5.2. Introduce the needle through the punctured shell at the injection spot and insert it all the way through, at a perpendicular angle, until the base of the needle touches the shell surface (Figure 2C).

NOTE: This positions the needle tip to deliver the syringe content into the allantoic cavity (in embryonated eggs) or the albumen (in non-embryonated eggs).

5.3. Carefully and slowly inject 100 µL of the inoculum.

NOTE: It is crucial to keep the angle of the needle as still as possible, when inside an embryonated egg, to avoid causing lethal injuries on the embryo.

5.4. Carefully and slowly remove the needle and cover the injection spot with a round sticker, to seal the inoculation entry site.

NOTE: For convenience and clarity, use stickers with a different color for each condition/group of eggs.

5.5. Return the inoculated eggs to the incubator. Incubate for 2 days.

NOTE: Eggs with developing embryos should not be out of the incubator (or at a temperature below 37.7 °C) for more than 30 min. Perform the inoculation in batches, if necessary.

### 6. Assessing the presence of culturable cells in the inoculation samples

6.1. Dispense 100 µL of VBNC bacterial suspension into multiple wells of a 96-well microplate. NOTE: Number of wells ≥ 2 × number of inoculated (embryonated + non-embryonated) eggs.

6.2. If using the control conditions mentioned in step 5.1, dispense 100 µL of each inoculum (i.e. mineral water or culturable bacteria) into at least three wells in the same 96-well microplate.

6.3. Using a multichannel pipette, add 100 µL of BHI medium to each of the wells prepared in step 6.2.

6.4. Incubate the plate at 37 °C under static conditions to promote bacterial growth in wells containing culturable bacteria.

NOTE: Depending on the presence of culturable bacteria and their regrowth rate in these conditions, one or more incubation days may be necessary to fully reveal wells containing culturable bacteria.

3.5. For each inoculum, count the number of wells scoring positive and negative for bacterial growth.

### 7. Monitoring embryo viability

7.1. The day after the inoculation, candle the embryonated eggs to check for embryo lethality.

7.2. Put aside eggs containing dead embryos, and discard them according to the institutional guidelines for infected biological waste disposal.

### 8. Processing of embryonated eggs

8.1. Recover the embryonated eggs from the incubator, and candle to check for embryo lethality.

NOTE: Dead embryos are discarded and therefore not considered in the final results.

8.2. Remove the sticker from the egg and disinfect the top end of the shell (covering the air pocket) with a tissue paper soaked with 70% (v/v) ethanol.

8.3. Using a clean pair of dissection scissors, cut the shell open from the injection spot to expose the air pocket. Using a clean pair of dissection tweezers, tear open the inner shell membrane that separates the air pocket from the rest of the egg.

8.4. Carefully empty the egg contents into a sterile Petri dish. Using a pair of tweezers, isolate and transfer the embryo onto a new Petri dish. Wash the embryo with sterile PBS, if necessary.

8.5. Transfer the embryo into a 15-mL centrifuge tube containing 4 mL of sterile PBS.

8.6. Dissociate the embryo using a homogenizer (speed: 10,000 rpm).

NOTE: Clean the tip of the dispersion tool between each homogenization by passing it sequentially in sterile PBS, 70% (v/v) ethanol, and again sterile PBS.

8.7. Plate 500 µL of the embryo homogenate on BHI agar. Incubate the plates at 37 °C overnight (or longer, if necessary).

NOTE: Optionally, serial dilutions of the homogenate can be plated in case elevated CFU numbers (i.e. >300) are expected in the undiluted homogenate.

8.8. For each inoculation group, count the number of eggs scoring positive and negative for *L. monocytogenes* growth on agar plates.

NOTE: Despite the aseptic conditions under which the embryo extraction and homogenization is performed, the use of non-selective agar medium for plating embryo homogenates entails a risk of contaminant growth. The use of a genus/species-specific growth medium or supplementation with a selective antibiotic (for which only the tested bacterial strain is resistant) can reduce or prevent such contamination.

### 9. Processing of non-embryonated eggs

9.1. Recover the non-embryonated eggs from the incubator. Repeat steps 8.2 and 8.3 to gain access to the egg albumen.

9.2. Using a micropipette, collect 500 µL of albumen and spread directly on BHI agar. Incubate the plates at 37 °C overnight or longer, if necessary.

9.3. For each inoculation group, count the number of eggs scoring positive for *L. monocytogenes* growth on agar plates.

## REPRESENTATIVE RESULTS

To test the potential of the chicken embryo model to revive cell wall-deficient VBNC forms of *L. monocytogenes* generated by starvation in mineral water, it was important to articulate the timing of preparation of the VBNC bacterial inoculum (≥28 days) with that of the embryonated eggs (6 days). Replicate suspensions of *L. monocytogenes* were thus set up at a concentration of 10^6^ CFU/mL in mineral water 28 days before the planned egg inoculation day. This starting bacterial concentration was shown to result in a desired residual culturability of <1 CFU/mL after 28 days^30^. A final verification of the bacterial cell culturability and viability levels of the suspensions resulted in the selection of a VBNC *L. monocytogenes* inoculum containing 10^6^ viable cells and 0.5 culturable cells per mL. One hundred microliters of this inoculum (i.e. 10^5^ viable cells and <0.1 CFU) were delivered into groups of embryonated and non-embryonated eggs. Control groups were prepared in parallel by injecting embryonated and non-embryonated eggs with the same dose volume of mineral water or of a 5×10^3^ CFU/mL suspension of *L. monocytogenes* grown overnight in BHI broth.

Two days post-inoculation, eggs were processed to assess the presence of culturable *L. monocytogenes*. As expected, no bacterial growth was recovered from the sterility control group eggs injected with mineral water, while 100% of eggs from the control group treated with culturable *L. monocytogenes* scored positive for growth on agar media (Table 1). Importantly, all embryos recovered from eggs inoculated with VBNC bacteria gave rise to *L. monocytogenes* growth on plate, contrasting with its complete absence on every plate spread with the content of a non-embryonated egg injected with the same VBNC bacteria (Table 1). This result underlines the requirement of the chicken embryo for the awakening process of VBNC *L. monocytogenes*, as previously reported^29^.

**Table 1:** VBNC *L. monocytogenes* revert back to a culturable state after passage in embryonated eggs. Embryonated and non-embryonated chicken eggs (6 days) were inoculated with 100 µL of mineral water only or with mineral water suspensions of VBNC (10^4^ cells, 28 days) or culturable (500 cells) *L. monocytogenes*. Two days later, embryos (or albumen in non-embryonated eggs) were recovered and plated on BHI agar to assess the presence of culturable *L. monocytogenes*. The frequency of culturable cells present in an inoculation dose before egg passage was determined by mixing 100 µL of each suspension with BHI broth in multiple wells of 96-well microplate, and scoring the number of wells with bacterial growth after incubation at 37 °C. Statistical significance was calculated using two-tailed Fisher’s exact test, with p-values ≥0,05 considered non-significant. This table was reproduced and adapted with permission from Carvalho et al.^30^

| Inoculum | Culturability in BHI before egg passage <sup>(1)</sup> | Culturability in BHI after egg passage <sup>(2)</sup> |  |  |  |
| --- | --- | --- | --- | --- | --- |
|  |  | Embryonated eggs | p-value <sup>(3)</sup> | Non-embryonated eggs | p-value <sup>(4)</sup> |
| Mineral water | 0/3 (0%) | 0/10 (0%) | >0.999 | 0/10 (0%) | >0.999 |
| VBNC *Lm* | 8/84 (9.52%) | 24/24 (100%) | 1.63×10<sup>-17</sup> | 0/18 (0%) | 0.344 |
| Vegetative *Lm* | 3/3 (100%) | 8/8 (100%) | >0.999 | 2/2 (100%) | >0.999 |

<sup>(1)</sup> Number of BHI wells with bacterial growth / Number of inoculated BHI wells
<sup>(2)</sup> Number of egg embryos with bacterial growth / Number of inoculated eggs
<sup>(3)</sup> Comparison of culturability before and after passage in embryonated eggs
<sup>(4)</sup> Comparison of culturability before and after passage in non-embryonated eggs

To determine the likelihood that the *L. monocytogenes* growth retrieved from embryonated eggs injected with VBNC bacteria comes from a true VBNC cell recovery, not from the regrowth of lingering culturable cells, it is necessary to compare the frequency of “inoculum doses” producing bacterial growth before and after egg passage. To obtain the former, multiple 100-µL doses of the VBNC bacterial inoculum were mixed with BHI, a nutrient-rich medium that does not support VBNC *L. monocytogenes* revival^31^, in a 96-well plate and incubated at 37 °C. A frequency of 9.5% of inoculated BHI wells positive for bacterial growth (8 in 84) differed significantly (Fisher’s exact test, *p*⍰=⍰1.6⍰×⍰10^−17^) from the 100% frequency of eggs showing recovered culturable bacteria (Table 1). This substantial and statistically significant difference strongly indicates that the *L. monocytogenes* growth recovered from embryonated eggs was largely due to reactivation of VBNC bacteria. In addition, unlike its vegetative forms, VBNC *L. monocytogenes* were unable to revive in non-embryonated eggs, supporting that VBNC cell revival in embryonated eggs is not due to the residual culturable bacteria in the inoculum.

## DISCUSSION

The public health and economical risks associated with “viable but non-culturable” (VBNC) forms of bacterial pathogens is a consequence of their capacity to evade detection by conventional microbial growth-based methods as well as of their tolerance/insensitivity to most antimicrobials used in clinical and food industry settings^25, 32^. It becomes therefore urgent to find research tools and models to better understand the mechanisms driving the transition, maintenance and revival of bacterial cells in a VBNC state.

In many aspects, the chicken embryo is an advantageous alternative to more conventional mammalian models of research (e.g. mouse, rat, rabbit). It is technically more practical to work with an oviparous organism, as its embryonic development is fast and occurs externally and independently of its progenitor. Importantly, the financial and logistical burdens associated with the production, housing and maintenance of chicken eggs and embryos are markedly lower. In addition, the easiness to obtain large numbers enables parallel testing of multiple experimental conditions and/or replicates and the generation of comprehensive and statistically robust data. Although non-mammalian models, such as fish or nematodes, can provide similar advantages, the chicken embryo – as a warm-blooded vertebrate – is evolutionarily closer to humans, and thus a better research model in areas such as immunology, disease modeling, development, and toxicology. Embryonic models are also simpler to work with, from an ethical standpoint, particularly during the first two-thirds of development, when a mature nociceptive system is not yet established^7^. This short time window is convenient for applications requiring quick turnaround times (e.g. drug testing^3^), but can pose restrictions for assays lasting into later embryogenesis stages or even post-hatching life (e.g. developmental and behavioral studies). The lack of an adaptive immune system also renders the chicken embryo a useful research tool in areas such as host-pathogen interactions and cancer biology^6^.

The chicken embryo has been extensively used in the study of microbial virulence^8–17^, but sparingly as a model to revive VBNC pathogens^26–29^. Here, we have provided a detailed methodological description of how this *in vivo* system can be successfully used for the rescue of VBNC *L. monocytogenes*.

A critical point to consider in this method is the issue of residual presence of culturable cells and their potential overpowering of true VBNC cell revival. The limitations of our VBNC cell production method to address this matter (VBNC cell “purity” levels are coupled with starting bacterial concentration) raise the need of employing alternative approaches to eliminate “contaminating” culturable cells altogether or reduce them to a negligible level. Such approaches may include, for instance, neutralization/killing via antimicrobials targeting exclusively culturable cells. Alternatively, physical removal of the culturable cell population can be achieved based on unique morphophysiological characteristics (e.g. cell shape, surface or cytoplasmic structures or molecules) via flow cytometry. This approach has the added advantage of allowing the isolation/purification of VBNC cell subpopulations with different ages (i.e. formed at different time points), which can be also tested *in ovo* for their revival capacity.

Other points of this method requiring attention include the careful handling of embryonated eggs, particularly during and immediately after the injection step (to avoid unintended needle-associated trauma or lethality), and striving to reduce the manipulation time outside of the optimal incubation temperature. In addition, despite the aseptic conditions under which the retrieval, processing and plating of embryos should be performed, unwanted microbial growth on agar plates may still occur. This can be solved by the use of agar media with a formulation (basal composition and/or supplements) selective for the bacterial species of interest, or by using a strain possessing resistance to a given antibiotic.

This work showcases the chicken embryo as a simple yet powerful and cost-effective *in vivo* model to investigate the mechanistic aspects of bacterial dormancy. In this context, it can potentially be used to identify bacterial factors implicated in the VBNC state awakening process, through the screening of individual bacterial mutants or on a genome-wide scale by employing a mixed library (e.g. transposon mutant library).

## ACKNOWLEDGEMENTS

This work was supported by a grant from Micalis Institute (AAP Micalis FAMe 2023). F.C. was supported by postdoctoral grant funding from Agence Nationale de Recherche (THOR: ANR-20-CE15-0008; PERMALI: ANR-20-CE35-0001).

